# Mapping multi-modal dynamic network activity during naturalistic music listening

**DOI:** 10.1101/2023.07.05.547865

**Authors:** Sarah EM Faber, Tanya Brown, Sarah Carpentier, Anthony Randal McIntosh

**Affiliations:** Institute for Neuroscience and Neurotechnology; The Virtual Brain; BEWorks

## Abstract

The human brain is a complex, adaptive system capable of parsing complex stimuli and generating complex behaviour. Understanding how to model and interpret the dynamic relationship between brain, behaviour, and the environment will provide vital information on how the brain responds to real-world stimuli, develops and ages, and adapts to pathology. Modelling together numerous streams of dynamic data, however, presents sizable methodological challenges. In this paper, we present a novel workflow and sample interpretation of a data set incorporating brain, behavioural, and stimulus data from a music listening study. We use hidden Markov modelling (HMM) to extract state timeseries from continuous high-dimensional EEG and stimulus data, estimate timeseries variables consistent with HMM from continuous low-dimensional behavioural data, and model the multi-modal data together using partial least squares (PLS). We offer a sample interpretation of the results, including a discussion on the limitations of the currently available tools, and discuss future directions for dynamic multi-modal analysis focusing on naturalistic behaviours.

## Background

In a letter to his son written in 1892, the painter Camille Pissarro wrote, “Everything is beautiful, all that matters is to be able to interpret.” Pissarro was referring to the work of his once-mentor, Jean-Baptiste-Camille Corot, but the crux of his sentiment, that to convey the breadth of the natural world, one must know what to attend to and how to render it in another form, is quite relevant to modern neuroscience. Understanding how the human brain operates in the real world has long been a goal of neuroscience research with computational frameworks emerging to capture more of how, and when, the brain operates. A flashpoint for several fields has been exploring the link between brain and behaviour to identify brain-based biomarkers capable of predicting behavioural outcomes and/or variability. This is particularly relevant in aging and health research where individual physiological outcomes may vary widely from predictions made using behavioural measures. From here, it would theoretically be possible to identify biomarkers for illnesses, predict treatment outcomes, and forecast who is more likely to exhibit resilience to injury and illness. The main challenges, however, are the methodological hurdles to analyzing high-dimensional brain and behavioural data, and interpreting meaningful findings from model output. In this work, we present a novel workflow and sample interpretation with the aim of showing how such an analysis can be conducted.

Foundational neuroimaging work has progressed from single region neural correlates to static networks, and now dynamic systems of networks and states in the growing field of dynamic functional connectivity. Certain frameworks in dynamic functional connectivity (e.g. hidden Markov modelling, leading eigenvector decomposition, sliding window analyses, Vidaurre et al., 2017; Cabral et al., 2017; Allen et al., 2014) allow for the exploration of regularly-occurring brain networks or states over time, gaining insight into how network activity emerges, develops, and decays over the course of behaviour (Misic & Sporns, 2016). Early network studies have been instrumental in identifying canonical networks related to specific cognitive states, such as the default mode network’s (DMN) involvement during wakeful rest, the salience network’s involvement in goal-directed tasks (Seeley et al., 2007; Sridharan et al., 2008), and dorsal and ventral attention networks’ roles in attention regulation (Corbetta & Shulman, 2002; Vossel et al., 2014).

Initially, these network studies existed largely in space rather than time with many studies collapsing functional brain activity over time to return static networks related to a particular behavioural state or task. While this work has provided a wealth of information on the brain’s network structure, the brain itself is a dynamic system and time-varying frameworks are required to understand how networks form and what drives shifts in network configuration (Calhoun et al., 2014; Hutchison et al., 2013). Estimating network dynamics has added much to our understanding of network functionality, including the extended role of the DMN. Initially thought to correspond to the brain at rest, through dynamic studies, it has been shown to be involved in numerous cognitive processes (see Spreng, 2012).

Through analysis of brain network dynamics, it is possible to uncover latent factors driving shifts in network configuration, but these latent factors do not necessarily come only from within the brain. The brain is a complex adaptive system that is connected, in space and in time, to other complex systems. A body, a history, the outside world – all of these elements contain information, weaving together across time to form complete experiences. Here, dynamic frameworks can be used to understand the shifting brain network landscape, and to model network timeseries data together with continuous behavioural or stimulus data to contextualize the brain network findings.

Music listening is uniquely suited to studying network dynamics. It is an information-rich signal stimulating numerous responses well-documented in prior behavioural and neuroimaging studies. Music listening work has identified neural correlates linked to music’s acoustic properties (Alluri et al., 2012), structural properties including timing and error detection (Burunat et al., 2014; Koelsch et al., 2007; Toiviainen et al., 2020), reward (Salimpoor et al., 2006; Gold et al., 2019), and emotional regulation (Carlson et al., 2015). Links between brain and behaviour have been identified through liking and familiarity with recent studies examining network responses to listening to familiar and preferred vs unfamiliar music in adolescents (Fasano et al., 2020) and younger and older adults (Belden et al., 2023, Faber et al., 2023). Music has also been employed to study physiological cues (see Mojtabavi et al., 2020), emotional regulation (Baltazar & Saarikallio, 2019; Baltazar et al., 2019), and social interaction through dance (Carlson et al., 2018; Carlson et al., 2019). These findings combined with its accessibility to and efficacy with clinical populations (Aalbers et al., 2017; Biasutti & Mangiacotti, 2018; Biasutti & Mangiacotti, 2021; Gold et al., 2004; Särkämö & Sihvonen, 2018) make music a compelling tool with which to investigate brain and behaviour via network and behavioural dynamics.

Quantifying continuous behaviour is not new in the field of cognitive psychology, but combining continuous behaviour tasks and neuroimaging is not broadly common. Exceptions are in two-alternative decision-making tasks (see Ratcliff, 1978; Ratcliff et al., 2016), musical improvisation (Da Mota et al., 2020), and movie watching (Whittingstall et al., 2010); but raise the issue of modality-specific event segmentation: how does one break up a continuous behaviour into units of meaningful information? Continuous behaviour can be rendered as a series of discrete steps, but embracing continuity can tell us more about how the brain behaves in the real world (Song & Nakayama, 2009).

One approach is applying dynamic frameworks from network studies to high dimensional behavioural and stimulus data. By transforming each data stream into a common conceptual space, it becomes possible to directly model the streams together while maintaining each modality’s salient characteristics. In this study, we demonstrate methodology for modelling brain, behaviour, and stimulus data together via a music listening study (Carpentier et al., 2020). We use hidden Markov modelling (HMM; Vidaurre et al., 2017) to estimate state timeseries data from the brain and the stimulus, manually estimate a state timeseries from the continuous behaviour task, and use partial least squares (PLS; McIntosh & Lobaugh, 2004) to uncover latent patterns between the modalities. While this work’s primary aim is to showcase the methodological workflow, we also offer an interpretation of the results and discuss future directions for multi-modal studies.

A note about the vocabulary and naming conventions: we use the terms “network” and “state” repeatedly throughout this work. We use the term “state” in the methodology and results sections to indicate these collections of brain regions are not consistent with canonical functional networks (such as DMN, salience network, etc). In the discussion, we use the term “network” consistent with the anatomical labels that best fit the states from the work of Uddin and colleagues (2019). In discussing prior work, we use “network” and “state” consistent with the original authors’ conventions.

## Methods

### Participants

Eighteen healthy right-handed adults (age range 19-35, mean age 26.0 years, 8 males) with normal hearing established via audiogram were recruited for a music listening task. All participants provided written informed consent as per the Baycrest Centre-University of Toronto Research Ethics Committee.

### Stimuli and apparatus

Stimuli were 40 excerpts of Western art music divided into perceptual (control) and emotional (experimental) tasks. Excerpts were between 40 seconds and two minutes long and were from the Renaissance to contemporary eras (Table 1). Excerpts were chosen to engender a range of subjective emotional reactions and were orchestrated in various styles, including solo instrumental, orchestral, chamber, choral, and operatic works. Vocal excerpts were in English, Italian, French, and German.

**Table 1:** List of stimuli for each task. Table adapted from “Complexity matching: brain signals mirror environment information patterns during music listening and reward” by Carpentier et al., 2020, Journal of Cognitive Neuroscience, 32 (4), p. 736.

| ***Emotional Songs*** |  |
| --- | --- |
| • Adams “Nixon in China, ‘Beginning’”<br>• Adams “Disappointment Lake”<br>• Bach “No. 3 Aria ‘Es Ist Vollbracht’”<br>• Barber “Adagio for Strings”<br>• Brahms “Intermezzo No. 2 in a Major, Op. 118”<br>• Delibes “Lakmé/Flower Duet”<br>• Elgar “Variation IX (Adagio) ‘Nimrod’”<br>• Galvany “Oh My Son”<br>• Gluck “Armide Act IV Air Sicilien”<br>• Goodall “Belief”<br>• Ives “Three Places in New England Orchestral Set No.1”<br>• Liszt “Totentanz”<br>• Monteverdi “Zefiro Torna”<br>• Mozart “Cosi fan tutte”<br>• Mozetich “The Passion of Angels”<br>• Part “Spiegel im Spiegel”<br>• Penderecki “Threnody to the Victims of Hiroshima”<br>• Puccini “O soave fanciulla” | • Rameau “Entrée de Polymnie”<br>• Richter “Vivaldi’s Summer”<br>• Rossini “Barbiere di Siviglia: Largo Al Factotum”<br>• Schroer “Field of Stars”<br>• Schumann-Liszt “Liebeslied (Widmung)”<br>• Staniland “Solstice Songs No. 2 Interlude”<br>• Stravinsky “Glorification of the Chosen One”<br>• Tarrega “Recuerdos De La Alhambra”<br>• Verdi “Messa Da Requiem: Dies Irae-Tuba Mirum Part 1”<br>• Verdi “Messa Da Requiem: Dies Irae-Tuba Mirum Part 2”<br>• Wagner “Die Walkurie, Act 3: Ride of the Valkyries”<br>• Wagner “Tristan Und Isolde/Act 2–Prelude” |
| ***Perceptual Songs*** |  |
| • Beethoven “Sonata in A Major Op. 69”<br>• Brahms “Violin Concerto in D, Op. 77-3”<br>• Glass “Glassworks Opening<br>• Haydn “Cello Concerto in D Major”<br>• Mozart “Symphony No. 40 in G-minor, K.550, Finale” | • Praetorious “Praeambulum”<br>• Strauss “Der Rosenkavalier Act III/Duet-Denouement and Grand Waltz-Coda”<br>• Strauss “September”<br>• Vivaldi “Concerto for Violin, Stings and Harpsichord in G”<br>• Vivaldi “Stabat Mater |

During music listening, participants completed a continuous ratings task. A two-dimensional cross was projected onto a computer screen and participants were instructed to track a cursor around the cross. In the control task, participants continuously tracked each excerpt’s perceived pitch and tempo with the axes of the cross corresponding to pitch (low-high) and tempo (low-high). In the experimental task, participants continuously reported their own inner emotional states with the axes of the cross corresponding to valence (unpleasant-pleasant) and arousal (relaxing-stimulating). The cursor reset to the centre of the cross at the beginning of each excerpt, and the ratings data were divided into five quadrants (Figure 1).

**Figure 1:**
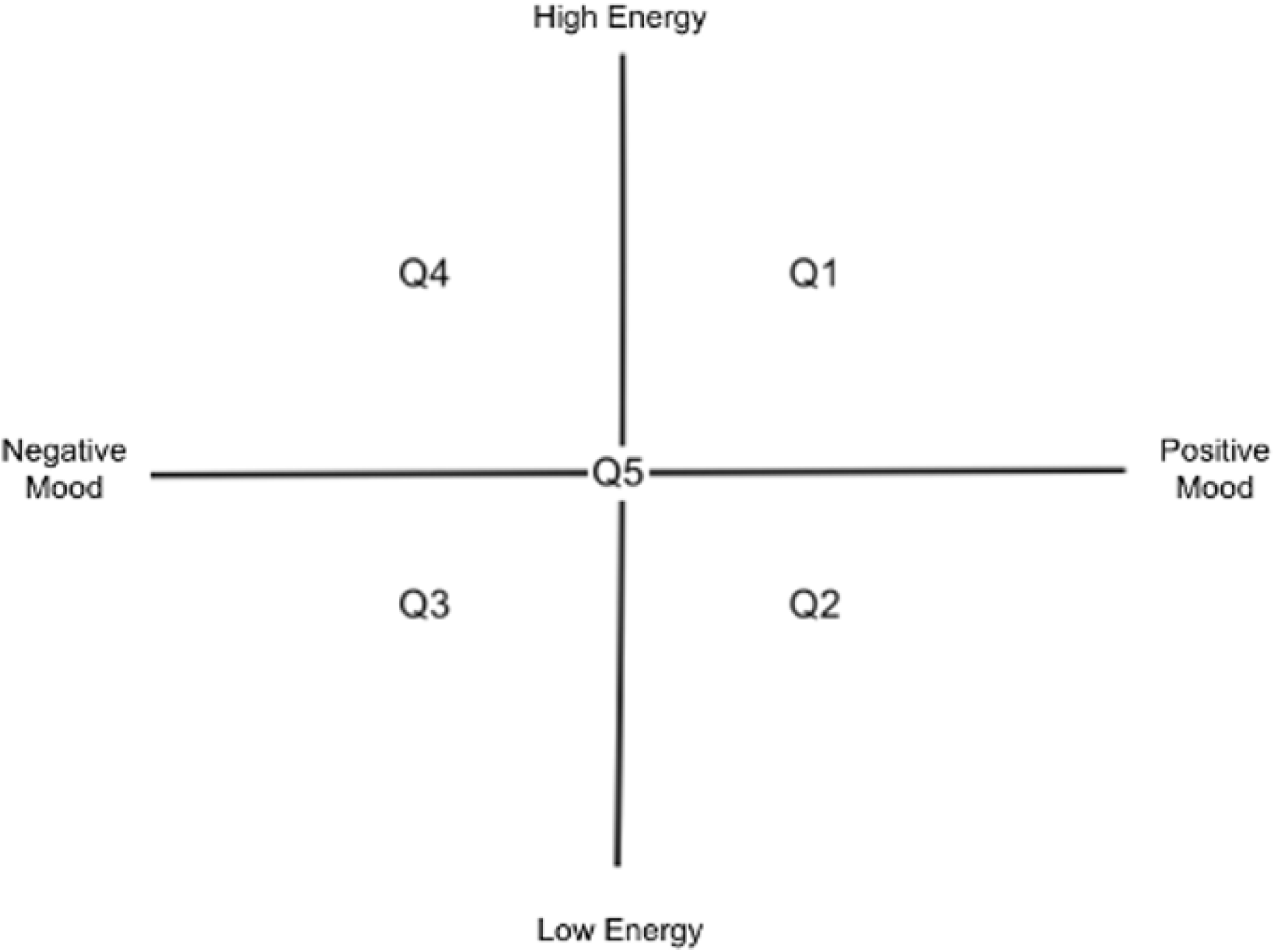
Mouse tracker cross with quadrant labels 1-5. The fifth “quadrant” is the centre starting position (x- and y-coordinates = 0,0)

### Procedure

Participants were provided orientation to the behavioural task apparatus and completed six practice trials. Participants completed five minutes of pre-task, eyes open resting state, then began the music listening tasks. Five control pieces were presented at the beginning of the task, followed by 30 experimental pieces. Five additional control pieces were presented at the end followed by five minutes of eyes open post-task resting state. Pieces were presented in one of three counterbalance orders: in the first, stimuli were ordered to “flow” from one piece to the next without sudden transitions. The second order was the reverse of the first, and the third order was random. Neither EEG nor behavioural measures differed significantly between the counterbalance orders, and the data were pooled for the remainder of the analysis.

### Data acquisition and pre-processing

Continuous EEG and behaviour data were collected simultaneously. EEG data were collected using a 64 + 10 channel Biosemi Active Two System with a sampling rate of 512 Hz. Data were imported to Brainstorm for pre-processing (Tadel et al., 2011). Raw EEG signals were bandpass filtered between 0.5 and 55 Hz to eliminate slow drift- and electrical line-related noise respectively, and re-referenced to the average of all electrodes. We used ICA for the detection of non-cerebral artifacts (ocular, muscular, and cardiovascular) for each participant. Data were epoched to 40 seconds (the length of the shortest music excerpt) and each epoch was visually inspected for noise related to loss of electrode connectivity. Data were source-localized using the Brainstorm implementation of sLORETA (see Tadel et al., 2011) projected into 68 regions of interest following the Desikan-Killiany atlas (Desikan et al., 2006; see Appendix 1 for a list of regions). Source waveforms were estimated for 15,768 equally-spaced vertices and the average current density of all vertices in each of the 68 regions of interest was calculated. This resulted in a 68-region*timepoint matrix for each task for each participant. These matrices were further concatenated into a single task*participant cell array with each cell corresponding to a single task (in this case, one music excerpt) for a single participant.

Of note, it is possible to complete these analyses with sensor-level data. We chose to proceed with source-localized data to facilitate the interpretation and comparison of our results with prior work, much of which is fMRI-based.

We imported the source-localized time series data to Matlab (R2016b), and estimated state properties using hidden Markov modelling (HMM) using the HMM-MAR toolbox (Vidaurre et al., 2016, Vidaurre et al., 2017). HMM is one of many algorithmic tools for the segmentation of high-dimensional time series’ into states that, for neuroimaging data, can describe spatial properties of the data. The inputs are not constrained to neuroimaging data, and the toolbox will work on any multivariate, high-dimensional data, including independent or principal component-decomposed data. The number of states are determined by the user *a priori* and the output includes a state time series (in the case of the HMM-MAR toolbox, the most likely sequence of states calculated using the Viterbi algorithm), state connectivity information (called *functional connectivity* in the HMM-MAR toolbox), and a transitional probability matrix describing the likelihood of transitions to and from each state. From these outputs, it is possible to calculate metrics describing the spatio-temporal properties of the timeseries.

In the present study, we ran HMM using Gaussian-distributed states. To maximize computational efficiency, we implemented HMM in a principal components analysis-reduced space preserving 90% of the variance of the raw data, and resampled the data to 256 Hz. To find an optimal value for K, we ran multiple HMM analyses with values for K between 5 and 20 (based on Vidaurre et al., 2017). We interrogated the transitional probability matrix from each estimation. The transitional probability matrix shows the pairwise likelihood of transitioning to and from each state, represented as a weighted, directed graph. Estimations with states that are over-represented (ie. always transitioned into) or under-represented (ie. rarely or never visited) can be considered to poorly fit the data. We found the estimations with a K of 8 and 12 fit the data best with a transitional probability matrix that did not feature states strongly over- or under-represented. We further completed initial analyses on these data using PLS, comparing the rest and music listening conditions. Both estimations returned one significant LV with a p value at or below 0.01 showing the contrast between the resting state and experiment tasks. Finding both estimations equal up to this point, we interrogated the spatial properties of the states. Each estimation featured states dominated by temporal regions, a comforting finding in a music listening study.

The estimation with a K of 12 featured bi- and uni-lateral temporal states, while the estimation with a K of 8 showed exclusively bilateral temporal states. Past findings have shown lateral effects in music feature processing (Alluri et al., 2012; Brown et al., 2006; Limb et al., 2006; Toiviainen et al., 2020), and we proceeded with the 12 state estimation.

We tested this estimation for stability by repeating the run 50 times and comparing the free energy values and state properties. The free energy values were broadly similar with a mean of 0.03 and a standard deviation of 0.0008. We further took the state matrices and ran STATIS, a way of interrogating the properties of symmetrical matrices (see Abdi et al., 2007), finding remarkably similar properties between states between model runs. This was encouraging, and we proceeded with the output from the estimation with a K of 12 for further analysis. We extracted the following metrics for each task block for each participant:

*State Visits*: how many times a state is visited during each task

*State Lifetimes*: how long, on average, a visit to a state lasts during a task

*Fractional occupancy*: the amount of time a participant spends in a state in total during each task

*Task-wise transitional probability matrices*: the pair-wise probability of transitions between states represented as a weighted, directed adjacency matrix

These metrics were averaged together by condition (rest, control task music listening, experiment task music listening). The resulting matrices were vectorized with the rows representing participants and columns representing HMM metrics, and used as input for partial least squares (PLS) analysis.

PLS is a multivariate analysis technique that describes the relationship(s) between matrices of measurements using singular value decomposition (SVD). Two common varieties are mean-centred PLS, which examines distributed activity differences between tasks, and behavioural PLS (also known as PLS correlation), which examines the multivariate correlations between two input matrices, often brain and behaviour. Both varieties return latent variables (LVs) that explain the relationships between matrices, accounting for progressively smaller amounts of covariance between matrices. Each LV includes a singular vector of weights for items within each input matrix, and a singular value, which is the covariance between the matrices (McIntosh et al., 1996). Statistical significance is established via permutation testing, and bootstrap estimation is used to determine the reliability of each LV (McIntosh & Lobaugh, 2004; Efron & Tibshirani, 1996). For this study, 500 permutations and 100 bootstrap estimations were used in each PLS analysis. No family-wise error correction was done for the number of PLS models, but the reliability of the model parameters assessed with bootstrap added another level of certainty to the statistical outcome. No family-wise error correction was implemented for the number of PLS models, but the reliability of the model parameters assessed with bootstrap estimations added another level of certainty to the statistical outcome.

The x- and y-position data from the mouse tracker were imported to Matlab (R2016b) and each timepoint was assigned to one of five mouse tracker states: quadrants one through four, and the starting neutral position. The advantage of HMM is that it allows for the exploration of patterns in high-dimensional data, but in low-dimensional data with a known quantity of non-temporal configurations, such as the present study’s behavioural data, states can be defined manually. This allowed us to calculate state fractional occupancy, lifetimes, and visits consistent with the brain HMM metrics. Because of the subjective nature of the task and the resetting of the cursor before each excerpt, we chose a simple quadrant-plus-neutral assignation for the behavioural data rather than further stratifying the quadrants.

We extracted features from the music data (Table 2) using the Music Information Retrieval Toolbox (MIRToolbox; Lartillot & Toiviainen, 2007) to further contextualize the brain and behaviour data. These features were extracted for each of the 40 pieces at 10 Hz. This sampling rate was chosen to capture information from both low- and high-level musical features.

**Table 2:** MIR features State estimation: Brain.

| *Feature* | *Description* |
| --- | --- |
| Zero Crossing | *Noisiness: how many times the signal crosses the 0 amplitude line* |
| Roughness | *Sensory dissonance from beating phenomenon* |
| Amplitude (root mean squared) | *Amplitude, a measure of loudness/energy* |
| Spectral Flux | *Distance between successive spectral frames* |
| Sub-band Flux 1-10 | *Same as above, decomposed into 10 sub-bands (ascending Hz)* |
| Pulse Clarity | *Estimate of beat regularity* |
| Mode | *Estimate of major or minor modality* |
| Key Clarity | *Strength associated with the most likely key(s)* |
| Novelty | *Probability of transitions between successive frames* |

Low-level features (zero-crossing, roughness, amplitude, and spectral and sub-band flux) describe acoustic/spectral properties of sound and are conveyed at rapid timescales. High-level features (pulse clarity, mode, key clarity, and novelty) describe properties of musical structure and are conveyed at longer timescales.

As with the EEG data, we completed multiple HMM estimations with varying values of K on the music data. We found a K of 5 to return a transitional probability matrix that did not feature over- or under-represented states. We tested this estimation for stability with a further 50 model runs and compared the free energy values and state properties. The mean free energy value across estimations was 0.048 with a standard deviation of 0.00077, indicating stability between estimations with the selected K. We again compared the state matrices using STATIS, finding a high degree of similarity between estimations. We proceeded with the K=5 estimation for further analysis.

The mean of each brain state extracted from HMM was calculated using the getMean function from the HMM-MAR toolbox (Vidaurre et al., 2017). This function projects the ROIs through each state’s principal components analysis (PCA) component scores, resulting in a sources*state matrix where positive and negative values refer to component loadings. State means were thresholded at the absolute 70th percentile and plotted onto a cortical surface using the ggseg package in R Studio (Mowinckel & Vidal-Piñeiro, 2020). The plots are displayed in Figure 2 and regions of interest and network labels are outlined in Table 3.

**Figure 2:**
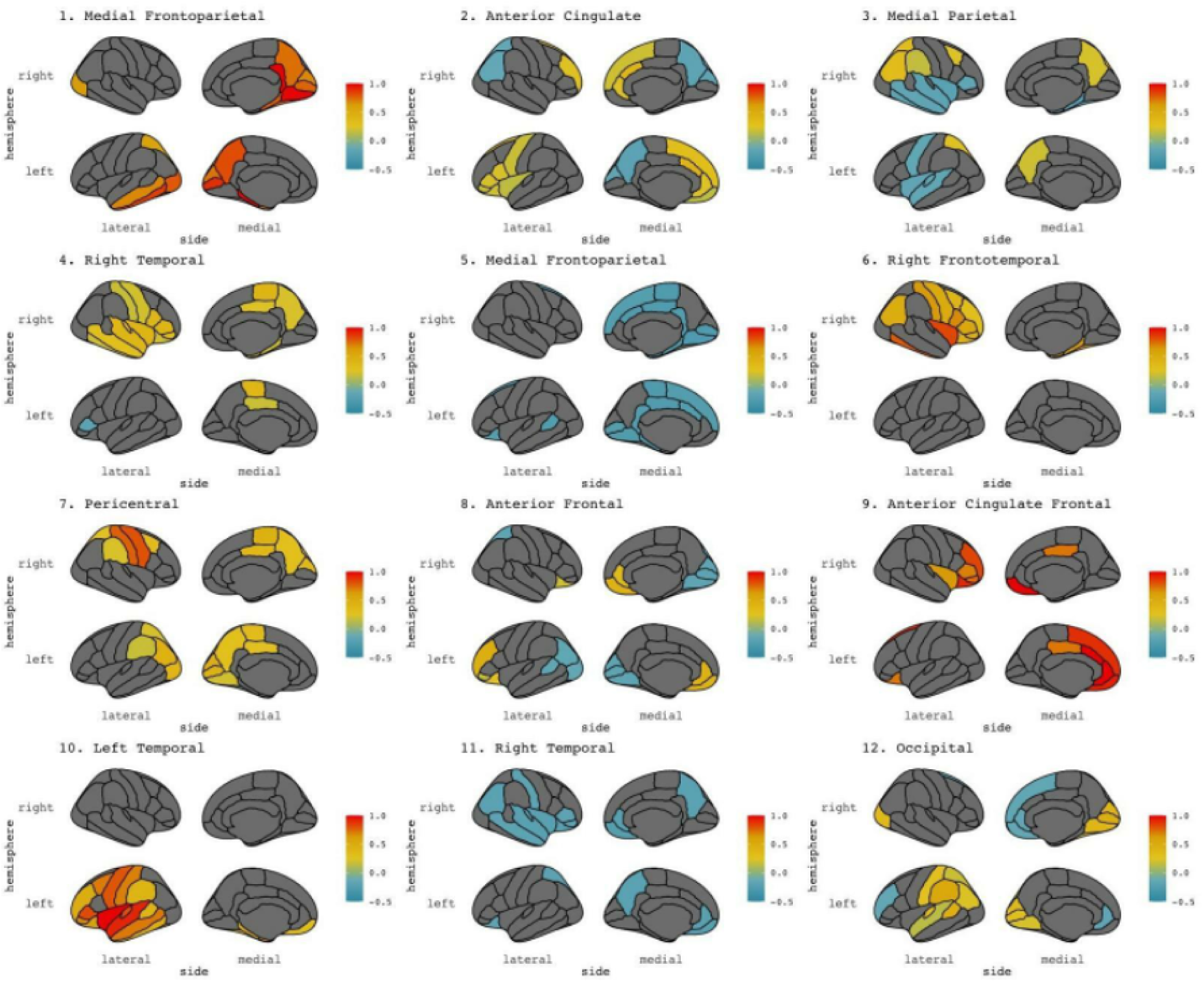
Mean state activity maps projected on cortical surfaces. Colours represent the voltage of each ROI relative to its loading in the PCA-reduced space from the HMM initialization. Warm colours correspond to a strong positive loading, cool colours represent a strong negative loading.

**Table 3:** Brain state regions of interest and network labels. Network labels are based on the work of Uddin et al. (2019).

| State | Main Regions | Network |
| --- | --- | --- |
| 1 | Activity in bilateral occipital cortices, bilateral precuneus, bilateral parahippocampal gyri, and left medial temporal lobe | Medial frontoparietal network |
| 2 | Activity in bilateral anterior cingulate cortex regions | Anterior cingulate network |
| 3 | Activity in bilateral precuneus and superior parietal, right precentral; lower relative activity in left precentral, bilateral temporal regions | Medial parietal network |
| 4 | Activity in left posterior cingulate and paracentral, right frontal, temporal, and parietal regions | Right temporal network |
| 5 | Activity in bilateral paracentral, cingulate, pericalcarine, and lingual regions | Medial frontoparietal network |
| 6 | Activity in right frontal, temporal, and parietal regions | Right frontotemporal network |
| 7 | Activity in bilateral parieto-occipital regions | Peri-central network |
| 8 | Activity in bilateral frontal regions, lower relative activity in bilateral parieto-occipital regions | Anterior frontal network |
| 9 | Activity in bilateral orbitofrontal regions, right inferior frontal gyrus regions | Anterior cingulate frontal network |
| 10 | Activity in left frontal, temporal, and parietal regions | Left temporal network |
| 11 | Activity in bilateral frontal, temporal, and insular regions | Right temporal network |
| 12 | Activity in left bank of superior temporal sulcus, left-side supramarginal gyrus, and occipital regions | Occipital network |

### State Estimation: Music

The MIR feature HMM returned five states, which we interpret as follows: State 1 corresponds to the acoustic texture of the sound, State 2 to prevalence in the mid-high frequency range, State 3 to prevalence in the low frequency range, State 4 to suddenness, and State 5 to predictability (Figure 3).

**Figure 3:**
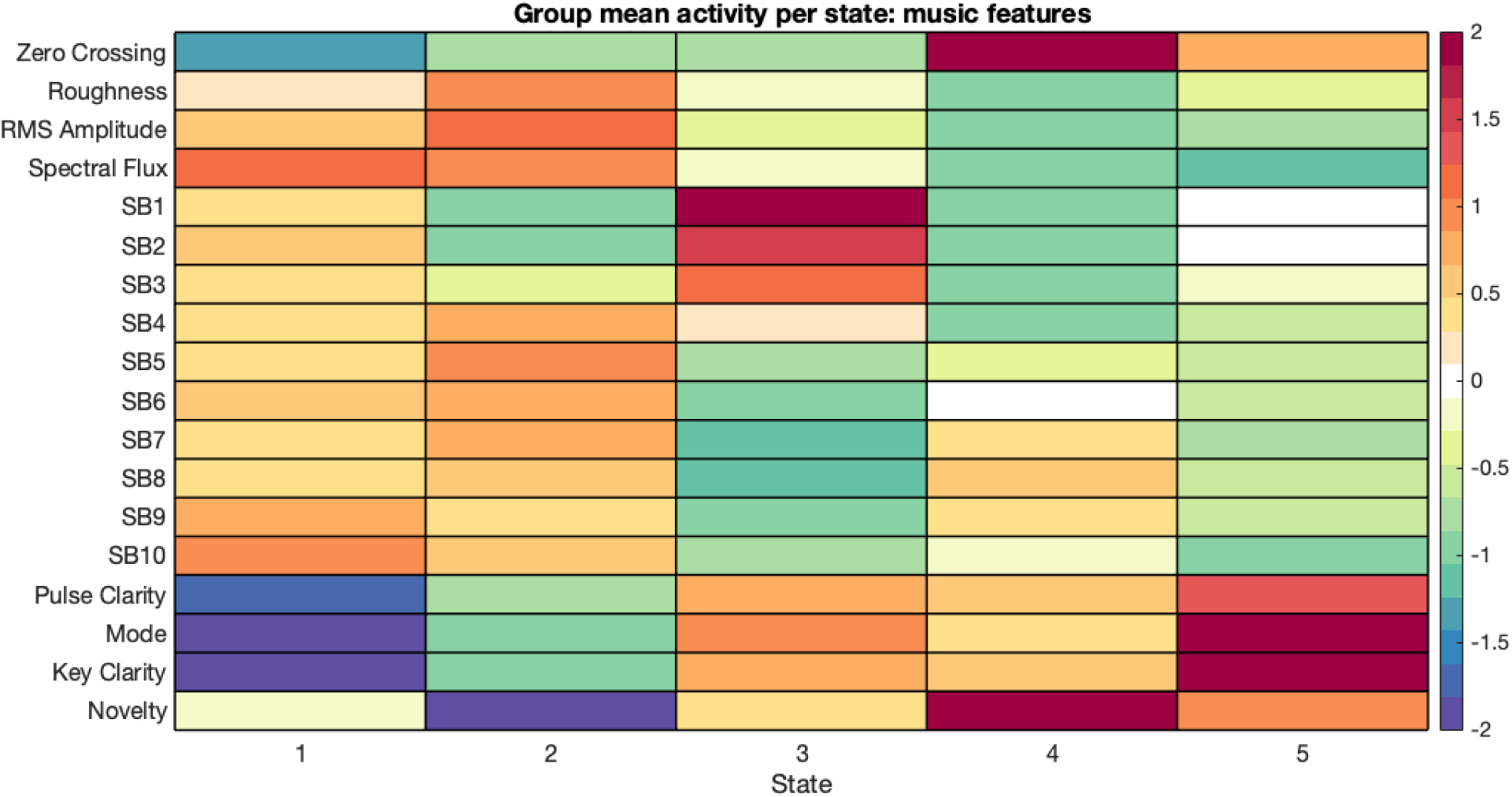
MIR feature states.

States 1 through 3 describe the acoustic properties of the sound. The musical information in these states is conveyed at rapid timescales and not dependent on prior information, making states 1-3 context-free. States 4 and 5 describe context-dependent structural properties of the sound. The musical information in these states is conveyed more gradually and is contingent on prior information. For clarity, we have grouped these states into two categories: *context-free features* and *context-dependent features*.

We extracted state fractional occupancy from the music HMM, but not lifetimes or visits due to the low sampling rate of the music data.

## Results

### Brain and behaviour: between-task analyses

We extracted state fractional occupancy, visits, and lifetimes from the EEG HMM estimation and analyzed the data for task differences between rest, control, and experimental conditions using mean-centred PLS. One significant LV was returned from the fractional occupancy and lifetime analyses, showing higher fractional occupancy in brain states 3, 4, and 8 during rest and higher fractional occupancy in state 5 during music listening in the experimental condition (*p* = .034).

Higher lifetimes were observed in states 1 and 5 during music listening in the experimental condition (*p* = .026). No significant LVs were returned from the visit analysis. Follow-up analyses were performed between the music listening tasks, returning no significant results (*p* = 0.14).

We next extracted rest- and task-wise transitional probability matrices and completed a mean-centred PLS analysis. No significant LVs were returned (*p* = 0.33). We ran a second mean-centred PLS on the transitional probability matrices between the two music listening conditions, which returned no significant LVs (*p* = 0.97).

### Brain and behaviour: within-task analyses

We next modelled the brain visit, lifetime, and fractional occupancy data together with the mouse tracker data for each music listening condition using behavioural PLS. No significant effects were returned at the individual task level (*p* = 0.10 for the control task, *p* = 0.38 for the experiment task). Combined behavioural PLS analyses were not performed as the mouse tracker quadrants correspond to features unique to each task (music perception and music-evoked emotions) and cannot be equated.

We completed behavioural PLS between transitional probability data and mouse tracker values within-task, which returned one significant LV (*p* = 0.01) showing an interaction between lifetimes in the low valence/low arousal quadrant and likelihood of transitioning to other states in the experiment condition (Figure 4).

**Figure 4:**
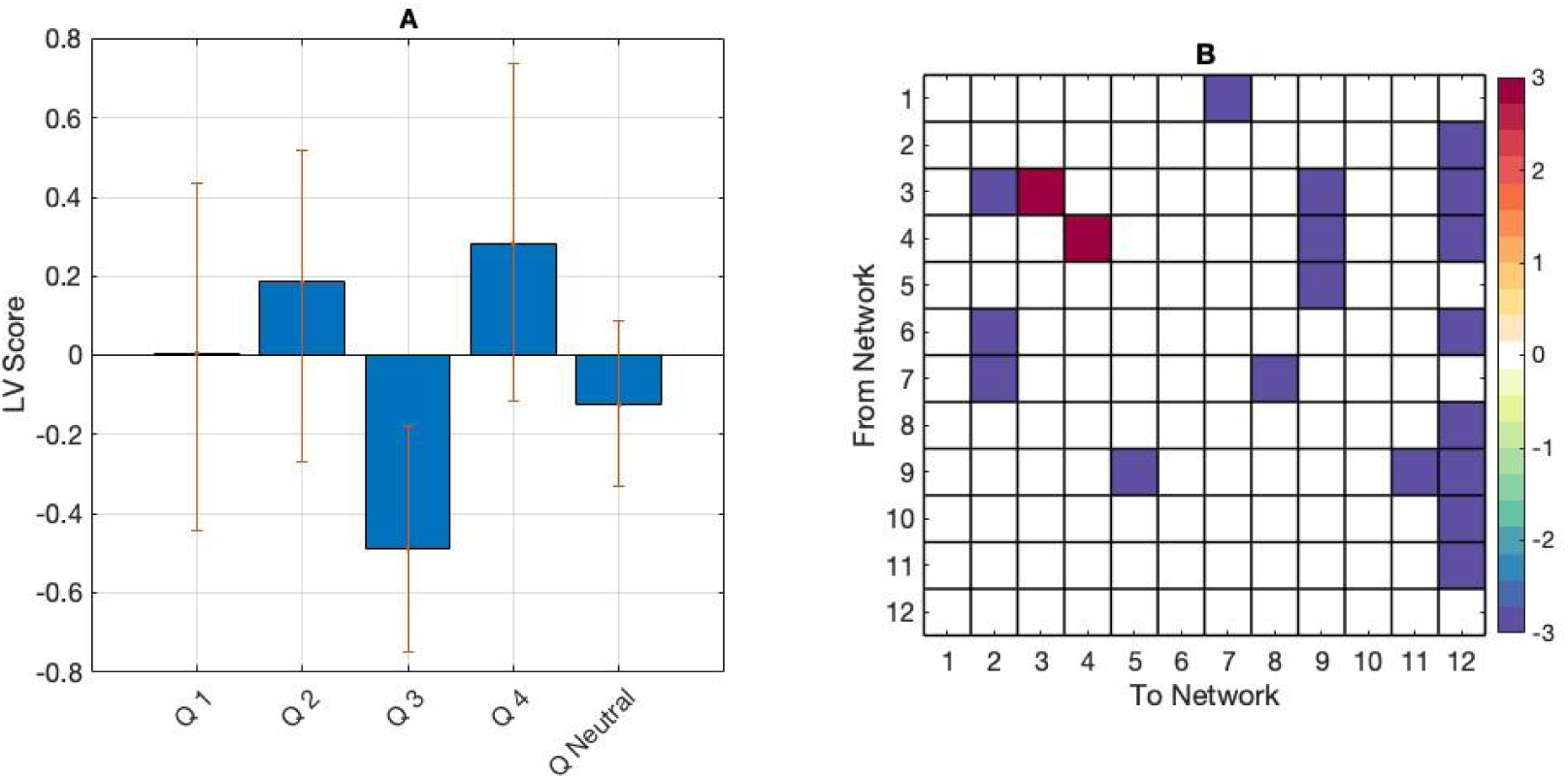
(A) PLS contrasts between lifetimes in mouse tracker quadrants (Q) 1-5. Error bars were calculated using bootstrap resampling and reflect the 95% confidence interval. The contrasts show an effect of behaviour on transitional probabilities (B) with a higher likelihood of transitioning to other states and a lower likelihood of staying in states 3 and 4 while reporting a low arousal/low valence emotional state.

### Brain and stimulus: between-task analyses

We introduced stimulus data into the analysis and correlated the EEG visits, lifetimes, and fractional occupancy with the fractional occupancy from the music HMM estimation. We ran mean-centred PLS on these correlation matrices, which returned one significant LV (*p* < .01) showing a task effect. In the control task, brain state visits and lifetimes were more positively correlated with MIR state fractional occupancy, particularly in brain states 4, 5, 7, 8, and 10. In the experiment task, MIR fractional occupancy was more positively correlated with brain fractional occupancy, particularly in brain states 3, 8, and 9 (Figure 5).

**Figure 5:**
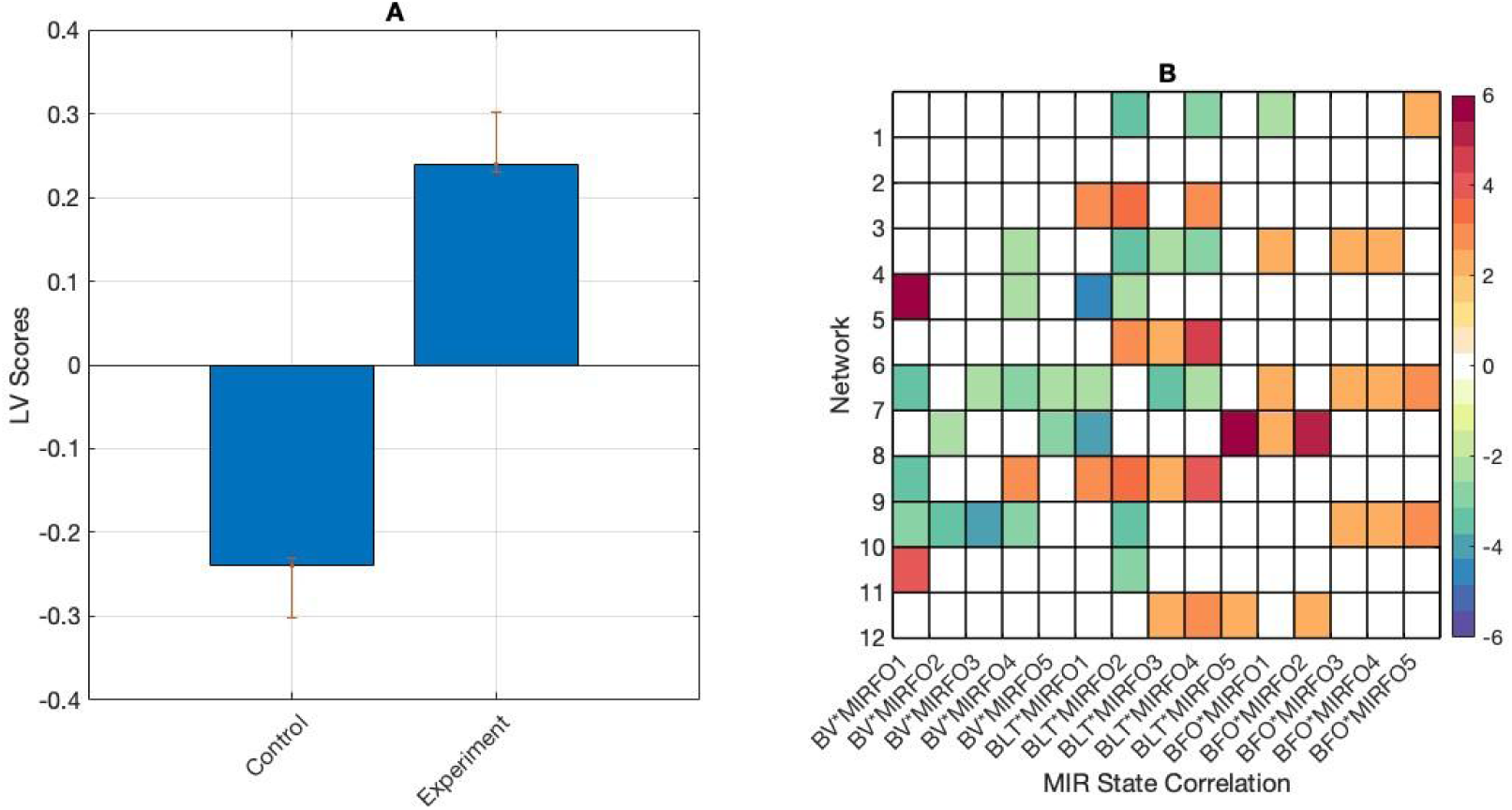
(panel A) PLS contrasts between tasks, and correlations between brain and stimulus measures. Error bars were calculated using bootstrap resampling and reflect the 95% confidence interval. The contrasts show an effect of task on correlations between brain and stimulus measures (panel B). Abbreviations are as follows: brain visits (BV), brain lifetimes (BLT), brain fractional occupancy (BFO), and MIR fractional occupancy (MIRFO).

We next correlated the MIR state fractional occupancy data with the brain transitional probability matrices. To do this, we vectorized the task-averaged transitional probability matrices and correlated them with the task-averaged MIR fractional occupancy. An initial between-task mean-centered PLS returned no significant LVs (*p* = 0.89). We subsequently correlated the piece-wise transitional probability matrices with the piece-wise MIR fractional occupancy and again found no significant differences between the tasks via mean-centred PLS (*p* = 0.78).

We then averaged the correlation matrices by MIR fractional occupancy category (context-free, and context-dependent) and ran a mean-centred PLS. This analysis returned one significant LV (*p* = .001) showing the contrast between transitional probabilities correlated with context-free and context-dependent MIR states in the control task only. Context-free MIR states were positively correlated with transitions involving brain states 3, 5, 6, 7, 9, and 12; while context-dependent MIR states were positively correlated with transitions involving brain states 2, 5, 10, and 11 (Figure 6).

**Figure 6:**
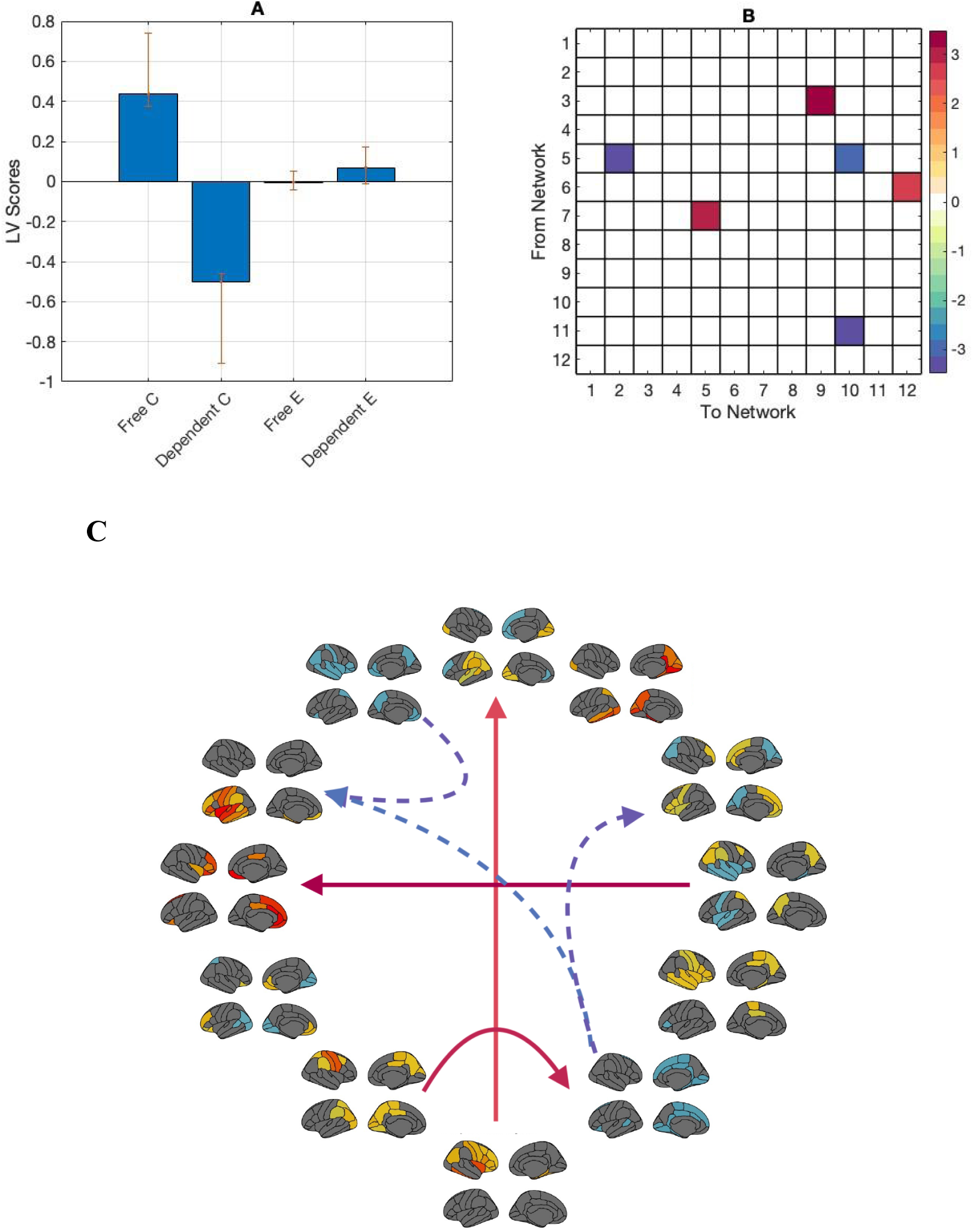
(panel A) PLS contrasts between context-free and context-dependent MIR states in the control task (Free C, Dependent C). Error bars were calculated using bootstrap resampling and reflect the 95% confidence interval. The contrasts show an effect of task and feature category in correlations between transitional probabilities and MIR features (panel B) . Positive correlations were present between numerous brain states and both context-free and context-dependent MIR states. Panel C shows the directed transitional probability correlations with the r-value *magnitude. Solid lines represent correlations with context-free MIR features, dashed lines represent correlations with context-dependent MIR features*.

### Behaviour and stimulus

To study the interactions between the stimulus and behaviour data, we correlated the mouse tracker data with the MIR state fractional occupancy data and ran a between-task mean-centred PLS. The analysis returned one significant LV (*p* <.01) showing a task effect in the correlations between behavioural ratings and MIR state features. In the control task, correlations were more strongly positive between fractional occupancy in MIR state 5 and fractional occupancy and lifetimes in mouse quadrant 1; between fractional occupancy in MIR states 1 and 3 and fractional occupancy and lifetimes in quadrant 2, and between fractional occupancy in MIR state 3 and fractional occupancy and lifetimes in quadrant 3. Control task correlations were more strongly negative compared to the experiment task between fractional occupancy in MIR states 1 and 3 and fractional occupancy and lifetimes in quadrant 1, fractional occupancy in MIR state 5 and fractional occupancy and lifetimes in quadrant 2, and fractional occupancy in MIR state 3 and fractional occupancy and lifetimes in quadrant 4 (Figure 7).

**Figure 7:**
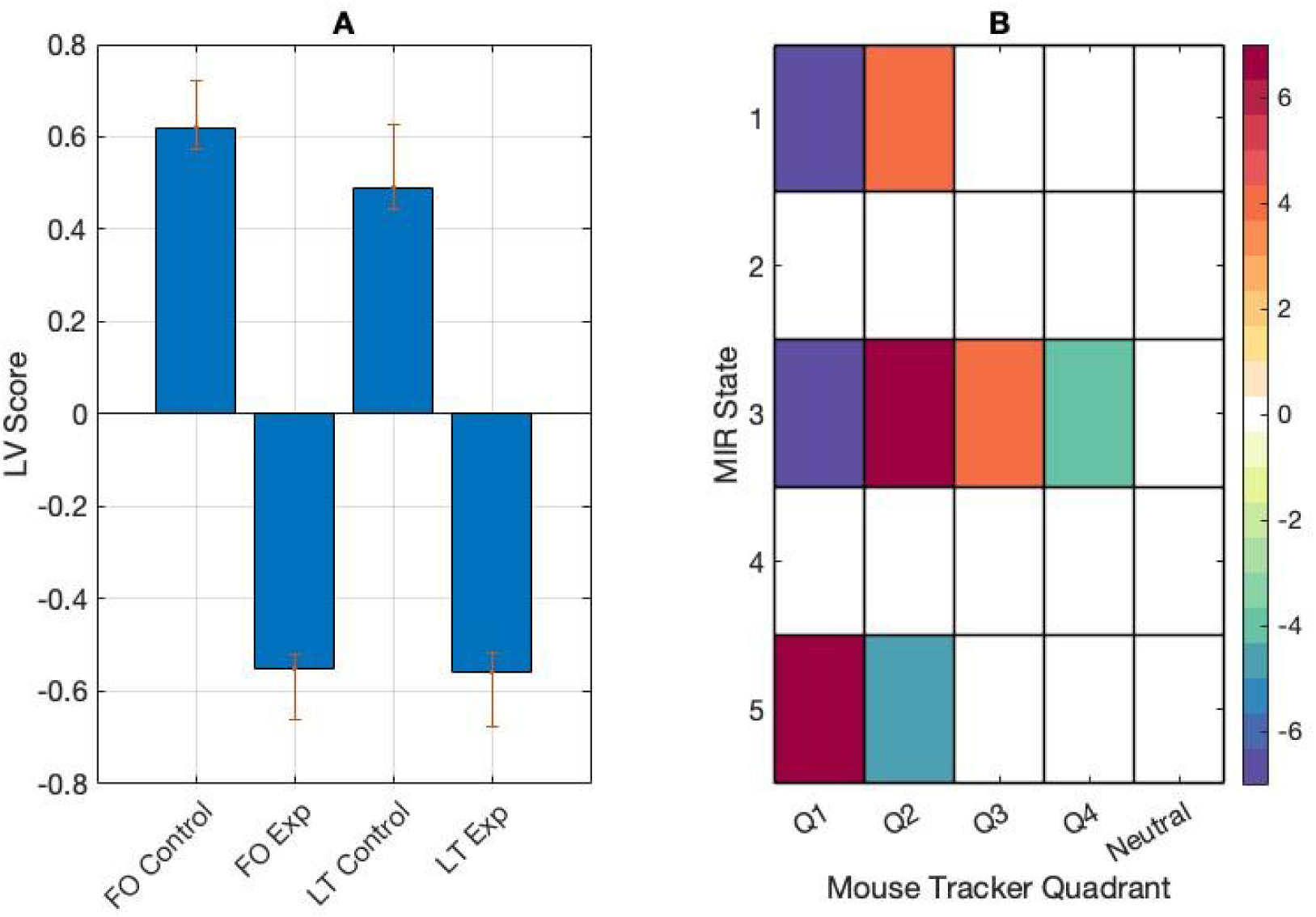
(panel A) PLS contrasts between fractional occupancy and lifetimes (FO and LT) in control and experiment (Exp) tasks, and correlations between behaviour and stimulus measures. Error bars were calculated using bootstrap resampling and reflect the 95% confidence interval. The contrasts show an effect of task on correlations between behaviour and stimulus measures in the mouse tracker quadrants (A) (panel B).

Follow-up mean-centred PLS analyses on averaged correlation data revealed task differences in one significant LV (*p* < 0.001). Correlations were more strongly positive between fractional occupancy in MIR state 2 and mouse tracker fractional occupancy and lifetimes in the control task. Correlations were more strongly negative between fractional occupancy in MIR state 1 and fractional occupancy and lifetimes in the control task, and more strongly positive between fractional occupancy in MIR state 1 and fractional occupancy and lifetimes in the experiment task (Figure 8).

**Figure 8:**
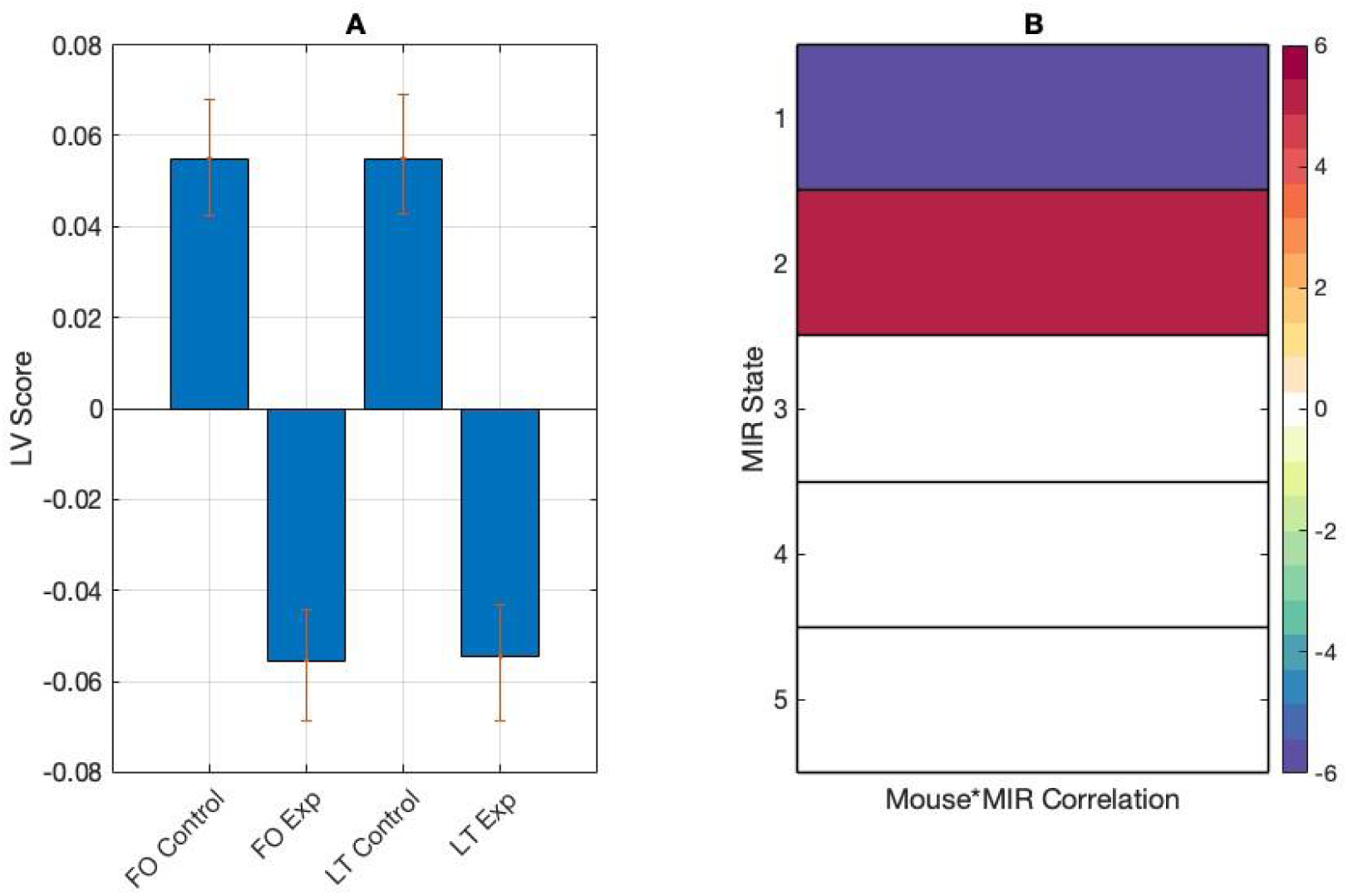
(panel A) PLS contrasts between fractional occupancy and lifetimes (FO and LT) in control and experiment (Exp) tasks and correlations between behaviour and stimulus measures. Error bars were calculated using bootstrap resampling and reflect the 95% confidence interval. The contrasts show an effect of task on correlations between behaviour and stimulus measures (panel B).

### Brain, behaviour, and stimulus

Finally, we ran a behavioural PLS between mouse tracker fractional occupancy and lifetimes in the starting neutral quadrant only (this quadrant was uniquely comparable between music listening tasks) and the correlations between MIR fractional occupancy and the brain transitional probability matrices. The analysis did not return any significant LVs (*p* = 0.26).

## Discussion

In this study, we presented an approach for modelling dynamic state timeseries data together with continuous behavioural and stimulus data using HMM and PLS. We estimated state timeseries for the high dimensional brain and music data using HMM, and manually estimated a state timeseries from the low dimensional behaviour data. We extracted measures describing the temporal properties of these timeseries’ and used PLS to identify latent patterns between each modality. These data were collected as a pilot study to test multi-modal recording in music listening. Numerous analyses we report here are non-significant, possibly owing to the small sample size and the subjective nature of the task. While the primary aim of this work is to demonstrate a methodological workflow, a brief interpretation of the results is included here.

Initial PLS analyses showed modality-specific fractional occupancy patterns between rest and the emotional music listening task. At rest, there was more overall occupancy in the medial parietal, right temporal, and anterior frontal networks; while in emotional music listening, there was more overall occupancy in the medial frontoparietal network. Analysis of the lifetimes, or average visit length, returned different results, with participants spending longer on average per visit in both medial frontoparietal networks while listening to music and attending to their emotions. These analyses demonstrate that HMM measures extracted from finely temporally resolved timeseries’ can be used to describe task-level differences.

The default mode network (DMN), consisting of many medial frontoparietal regions (Uddin et al., 2019), has shown higher levels of activity while listening to music that has been rated as liked (Wilkins et al., 2014; Pereira et al., 2011) and rated as more sad-sounding (Taruffi et al., 2017). These findings support DMN involvement in music listening with the added layer of complexity introduced by the continuous behavioural task.

Testing between the control and experiment music listening tasks returned no significant differences for either the HMM-extracted metrics or transitional probability matrices. This is not entirely surprising - both tasks, at the surface level, are quite similar: participants are listening to music and providing continuous ratings throughout each excerpt. The main difference is in the ratings themselves. In the perceptual control task, participants were asked to rate perceptual features in the musical signal, while in the emotional experiment task, they were asked to rate music-induced changes to their own emotional state. As such, both tasks were inherently subjective. Participants were not given training on perceptual feature rating, and as a result, perceptual ratings varied between participants (Carpentier et al., 2020).

The emotional data likewise varied between participants, and we did not initially expect to see a group-level effect between ratings data and brain metrics. One effect emerged in the experiment task, however. Overall time spent in the low arousal/low valence quadrant (analogous to feeling sad) was significantly positively correlated with the likelihood of transitioning between states, especially into networks involving the anterior cingulate, frontal, and occipital regions (Figure 4). These regions are linked to emotion processing and attention (Uddin et al., 2019), and visual processing, indicating a community of networks involved in attending to and reporting the target emotion during music listening. Time spent in this state was negatively correlated with the likelihood of staying in the medial parietal and right temporal networks. Regions in the medial parietal cortex have been shown to be active as part of the default mode network, while right-lateral temporal regions have been implicated in music processing, especially melody (Alluri et al., 2012). During reported low-valence, low-arousal mood, these networks are less likely to persist, indicating that default mode and auditory processing networks are less continuously occupied during interoceptive monitoring in this reported emotional state.

The emotional condition of sadness, represented in the low-arousal, low-valence quadrant is interesting here. Past work has shown that sadness is more consistent between individuals in a music-based emotion identification task (Faber & Fiveash, 2014), and sad-sounding music can stimulate participants to engage in more introspective behaviours (Taruffi et al., 2017).

Sad-sounding music does not necessarily herald the induction of a sad mood (Garrido et al., 2011), but the consistency in identifying sad music and the accompanying introspection may feed into the experience of sadness, offering a possible explanation as to why this quadrant is the lone group-level effect present.

Group-level effects emerged in the control task when we introduced the stimulus data into the analysis (Figure 5). More robust correlations were observed between brain visits and lifetimes between music features and the right temporal, medial frontoparietal, pericentral, anterior frontal, and left temporal networks. These networks are associated with auditory processing of melodic and rhythmic information (Brown et al., 2006; Alluri et al., 2012) and executive function (Toiviainen et al., 2020). In the control task, participants were asked to listen to music and make continuous judgements on perceptual features in the music. That this task employs bilateral temporal and anterior networks is pleasingly intuitive.

In the emotional listening experiment task, more reliable correlations were present between music features and overall occupancy in the medial parietal, anterior frontal, and anterior cingulate networks. These networks are more implicated in executive function and emotion processing than auditory processing (Uddin et al., 2019). These results suggest that, in the emotional listening task, fine-grained auditory processing may be of secondary importance to activity in interoceptive monitoring networks. These networks also track musical features, perhaps to continually validate the reported emotional state for continued accuracy.

When we correlated brain network measures with the MIR feature states, the perceptual task data showed two distinct between-network connectivity patterns: one correlated with context-free music features comprising numerous medial parietal, frontal, and cingulate networks; and the other correlated with context-dependent music features comprising medial, anterior cingulate, and bilateral temporal networks (Figure 6). Interestingly, involvement of the left temporal network is unique to the context-dependent circuit. Tempo is heavily context-dependent, and regions in the left temporal cortex have been previously linked to rhythmic perception specifically (Limb et al., 2006).

The emotional listening experiment task correlations did not return any significant results in this analysis, possibly due to the more internally driven nature of the task. With perceptual listening, participants are, hopefully, listening carefully to the music, which should be reflected in correlations between the timecourse of the musical signal and the timecourse of the brain data. In emotional listening, participants are listening to the music, but attending to their internal states, which may be influenced by the music, but do not follow the timecourse of the music in the same way perception will. Also, the induction of emotional states by music operates on a longer timescale which may only be correlated with the musical signal at coarser windows (Eich & Metcalfe, 1989; Västfjäll, 2002). When modelling the music excerpt and behaviour data together, we again found more correlations in the perceptual task data, particularly in context-free music states, supporting the idea that participants are more in sync with the stimulus in the perceptual task (Figure 7).

We did not find any significant patterns when modelling together brain, behaviour, and stimulus data, however, this analysis provides a useful starting point for future work.

In sum, we show that between-state connectivity patterns are correlated with continuous behavioural reporting and continuous music features during music listening, and vary depending on task demands. Prior work has explored state occupancy and connectivity in task-free music listening in fMRI (Faber et al., 2023, Faber et al., 2024), and future work would be well poised to merge the temporal richness of EEG and music feature data with a more open-ended listening task.

These results offer a framework for extracting network timeseries from continuous brain, behaviour, and stimulus data; and extracting interpretable interactions between the data streams using latent variable analysis. While many tools and frameworks for timeseries segmentation exist, we present a way to transform continuous behavioural and stimulus data into the same conceptual space as the brain data by using the same or similar network estimation techniques.

As we demonstrate, network estimation may be applied to any high-dimensional timeseries using methods like HMM, or manually estimated from low-dimensional timeseries’ in cases where states or networks can be readily identified or inferred. We also show that network prevalence and the transitional patterns between them change when modelled together with stimulus and behavioural data. Understanding how these streams are connected and how to model them will become vital as we strive to understand the workings of the brain under more naturalistic conditions. Much work is being done in dyadic and social interaction studies combining motion capture, eye tracking, and EEG recording (Koul et al., 2023); modalities that are amenable to the methods we present here. Future work combining brain and physiological data can use this workflow to investigate interactions and integrations between brain, body, stimulus, and others.

A consideration is the number of currently available tools for state and manifold estimation, including leading eigenvector decomposition (LEiDA, Cabral et al., 2017), UMAP (McInnes et al., 2018), TDA-Mapper (Saggar et al., 2022), and DyNeMo (Gohil et al., 2022). Each have unique strengths and while a thorough review is beyond the scope of this work, we would be remiss not to acknowledge them. This work used HMM as we found it possible to interpret and model the output across our three data streams, however it is entirely possible that another tool could be used. These methods are not without limitations (for two excellent reviews, see Gonzales-Castillo & Bandettini, 2018; and Lurie et al., 2020), but their potential is too intriguing to ignore. HMM and other semi-supervised algorithms, such as LEiDA (Cabral et al., 2017) rely on the user to determine an optimal number of states (K) *a priori*. There is no current consensus on determining an optimal value for K, leading to ambiguity and variability among studies using these methods. We have presented our method for K selection in ambiguous cases here and previously (Faber et al., 2023) in an effort to provide clarity. However, we acknowledge that high-dimensional data are inherently study-specific. These methods are gaining popularity, and standards will emerge with further study.

Another limitation is replicability when an algorithm relies on semi-random initialization like HMM and LEiDA do. Here, we consider the need to find ways to capture variability at the individual and task level if we expect to expand our understanding of how the brain operates under real-world conditions. Network studies using canonical regions or seeds selected *a priori* (see Bressler & Menon, 2010) tell us much about the most common or seemingly necessary components in functional networks; however, they cannot explain signal propagation to unexpected regions or transitions through networks that contain regions from both networks of origin and destination. Consider as well the limitations of what we consider to be canonical networks. Calls for a more anatomical taxonomy have emerged in recent years (Uddin et al., 2019) and raise vital points about how we evaluate functional labeling in network neuroscience. When the task or the participant population is under-explored, as is frequently the case in clinical research, canonical functional regions may not be the best fit for the data. Indeed, in the present study, initial exploration into nodes related to auditory networks returned a coarse, bilateral auditory network that does not reflect the nuance of music listening, its lateral components, nor its links to regions and networks outside the temporal lobes.

As research continues to investigate more complex questions and as technology evolves to capture and process greater volumes of data; researchers must consider adopting new methods capable of capturing the richness and variability of high-dimensional data. This work represents an authentic look at quantifying and interpreting naturalistic behaviour and provides a viable method for navigating through high-dimensional datasets. Future directions could also include constructing participant-level models and comparing between-participants for cohort- and/or population-level differences (the STATIS method (Abdi et al., 2005) may be particularly useful here, see Baracchini et al., 2023), adding another dimension to existing work on the link between brain and behavior. These methods are new, unfamiliar, and have great potential, particularly for those interested in understanding the brain as part of a larger interconnected system. In newness, we will encounter much uncertainty and with it, the opportunity to uncover different ways of interpreting the brain.

## Acknowledgements

We wish to express our sincere thanks to Giulia Baracchini for her comments on an earlier draft of this manuscript, and to Simon Dobri. Our thanks as well to the participants for their time and effort.

## Code and data availability statement

The code for all analyses detailed for this paper is available on the following GitHub page: https://github.com/fabsarah/HMM-PC-Study The data for this study are confidential, however, those interested in collaborating with the authors on future projects using this dataset with relevant data sharing agreements are invited to contact the corresponding author.

---

**Appendix 1:** Regions from the EEG data parcellation using the Desikan-Killiany atlas (Desikan et al., 2006)

| **Region** |  |
| --- | --- |
| caudal middle frontal L | caudal middle frontal R |
| entorhinal L | entorhinal R |
| frontal pole L | frontal pole R |
| lateral orbitofrontal L | lateral orbitofrontal R |
| medial orbitofrontal L | medial orbitofrontal R |
| pars opercularis L | pars opercularis R |
| pars orbitalis L | pars orbitalis R |
| pars triangularis L | pars triangularis R |
| rostral middle frontal L | rostral middle frontal R |
| superior frontal L | superior frontal R |
| insula L | insula R |
| paracentral L | paracentral R |
| banks superior temporal sulcus L | banks superior temporal sulcus R |
| inferior temporal L | inferior temporal R |
| middle temporal L | middle temporal R |
| parahippocampal L | parahippocampal R |
| superior temporal L | superior temporal R |
| temporal pole L | temporal pole R |
| transverse temporal L | transverse temporal R |
| caudal anterior cingulate L | caudal anterior cingulate R |
| isthmus cingulate L | isthmus cingulate R |
| posterior cingulate L | posterior cingulate R |
| rostral anterior cingulate L | rostral anterior cingulate R |
| fusiform L | fusiform R |
| inferior parietal L | inferior parietal R |
| postcentral L | postcentral R |
| precentral L | precentral R |
| precuneus L | precuneus R |
| superior parietal L | superior parietal R |
| supramarginal L | supramarginal R |
| cuneus L | cuneus R |
| lateral occipital L | lateral occipital R |
| lingual L | lingual R |
| pericalcarine L | pericalcarine R |

